# Origins of cell-to-cell variability, kinetic proof-reading and the robustness of MAPK signal transduction

**DOI:** 10.1101/021790

**Authors:** Sarah Filippi, Chris P. Barnes, Paul Kirk, Takamasa Kudo, Siobhan McMahon, Takaho Tsuchiya, Takumi Wada, Shinya Kuroda, Michael P.H. Stumpf

**Affiliations:** Centre for Integrative Systems Biology and Bioinformatics, Imperial College London, London SW7 2AZ, UK; Department of Cell and Developmental Biology, University College London, London WC1E 6BT, UK; Department of Genetics, Evolution and the Environment, University College London, London WC1E 6BT, UK; Department of Biophysics and Biochemistry, Graduate School of Science, University of Tokyo, Tokyo 113-8654, Japan; CREST, Japan Science and Technology Agency, Bunkyo-ku, Tokyo 113-0033, Japan; Institute of Chemical Biology, Imperial College London, London SW7 2AZ, UK

## Abstract

Cellular signalling processes can exhibit pronounced cell-to-cell variability in genetically identical cells. This affects how individual cells respond differentially to the same environmental stimulus. However, the origins of cell-to-cell variability in cellular signalling systems remain poorly understood. Here we measure the temporal evolution of phosphorylated MEK and ERK dynamics across populations of cells and quantify the levels of population heterogeneity over time using high-throughput image cytometry. We use a statistical modelling framework to show that upstream noise is the dominant factor causing cell-to-cell variability in ERK phosphorylation, rather than stochasticity in the phosphorylation/dephosphorylation of ERK. In particular, the cell-to-cell variability during sustained phosphorylation stems from random fluctuations in the background upstream signalling processes, while during transient phosphorylation, the heterogeneity is primarily due to noise in the intensity of the upstream signal(s). We show that the core MEK/ERK system uses kinetic proof-reading to faithfully and robustly transmits these variable inputs. The MAPK cascade thus propagates cell-to-cell variability at the population level, rather than attenuating or increasing it.

## Introduction

The behaviour of eukaryotic cells is determined by the intricate interplay between signalling and regulatory processes. Within a cell each single molecular reaction occurs randomly (stochastically) and the expression levels of molecules can vary considerably in individual cells (Bowsher and Swain, 2012b). These non-genetic differences can and frequently will add up to macroscopically observable phenotypic variation (Balázsi et al., 2011; Spencer et al., 2009; Spiller et al., 2010). Such variability can have organism-wide consequences, especially when small differences in the initial cell populations are amplified among their progeny (Pujadas and Feinberg, 2012; Quaranta and Garbett, 2010). Cancer is the canonical example of a disease caused by a sequence of chance events that may be the result of amplifying physiological background levels of cell-to-cell variability.

Better understanding of the molecular mechanisms behind the initiation, enhancement, attenuation and control of this cellular heterogeneity should help us to address a host of fundamental questions in cell biology and experimental and regenerative medicine. Characterisations of the origins of cell-to-cell variability in biological systems have so far generally related to gene expression (Elowitz et al., 2002; Hilfinger and Paulsson, 2011; Swain et al., 2002). However, cell-to-cell variability can arise without substantial contributions from transcriptional and translational processes and also characterizes signal transduction at the single cell level (Colman-Lerner et al., 2005; Jeschke et al., 2013). It has now become possible to track populations of eukaryotic cells at single cell resolution over time and measure the changes in the abundances of proteins. For example, rich temporal behaviour of p53 (Batchelor et al., 2011; Geva-Zatorsky et al., 2006) and Nf–*κ*b (Ashall et al., 2009; Nelson et al., 2004; Paszek et al., 2010) has been characterized in single-cell time-lapse imaging studies. But because these studies tracked a small number of cells continuously it is difficult to gauge the causes of cell-to-cell variability, due to lack of statistical power, but even this is now becoming possible (Selimkhanov et al., 2014). Alternatively, measurements can be obtained by quantitative flow or image cytometry (Ozaki et al., 2010) where data are obtained at discrete time points but encompass thousands of cells, which allows one to investigate the causes of cell-to-cell variability. In the present study, this latter methodology is applied to mitogen activated protein kinase (MAPK) signalling cascades. MAPK mediated signaling affects cell fate decision making processes (proliferation, differentiation, apoptosis and cell stasis) and cell motility. The mechanisms of MAPK cascades and their role in cellular information processing have been investigated extensively (Aoki et al., 2011; Kiel and Serrano, 2009; Mody et al., 2009; Piala et al., 2014; Sturm et al., 2010; Takahashi et al., 2010; Voliotis et al., 2014). Our aim is to gauge and characterize sources and effects of variability in MAPK signalling, focusing on the extracellular-signal-regulated kinase (ERK) and its response to external stimulation (see Figure 2A). We use quantitative image cytometry to probe the cellular abundancies of active ERK and its cognate kinase MEK in a large number of PC12 cells collected at different times. This is coupled to a detailed Bayesian analysis of mathematical models of the MEK-ERK signalling cascade, where we infer the modes of ERK phosphorylation and dephosphorylation and quantify the temporal effects of different sources of noise on the level of population heterogeneity in activated ERK and MEK.

The molecular causes underlying population heterogeneity are only poorly understood, but two notions have come to dominate the literature: intrinsic and extrinsic causes of cell-to-cell variability (Bowsher and Swain, 2012a; Hilfinger and Paulsson, 2011; Komorowski et al., 2010; Swain et al., 2002; Toni and Tidor, 2013) (see Figure 1A-D). The former refers to the chance events governing the molecular collisions in biochemical reactions. Each reaction occurs at a random time leading to stochastic differences between cells over time. The latter subsumes all those aspects of the system which are not explicitly modelled. This includes the impact of stochastic dynamics in any components upstream and/or downstream of the biological system of interest which may be caused, for example, by the stage of the cell cycle (which will affect cell sizes, transcription activity, and availability of free ribosomes and proteasomes) and the multitude of factors deriving from it. To date the most comprehensive characterisations of the interplay between intrinsic and extrinsic noise in biological systems generally relate to gene expression (Elowitz et al., 2002; Swain et al., 2002), using, for example, dual reporter assays which explicitly separate out extrinsic and intrinsic sources of variability (Hilfinger and Paulsson, 2011). These assays, however, cannot be used in all conditions; it is therefore important to develop alternative approaches that can distinguish between the different noise sources. Here we develop an *in silico* statistical model selection (Kirk et al., 2013) framework for this purpose and we demonstrate that we can confidently implicate extrinsic noise as the dominant factor giving rise to cell-to-cell variability in MAPK signalling. Further analysis of the MAPK dynamics allows us to highlight and attribute the origins of the extrinsic variability to biological processes upstream. In particular, we will show that the cell-to-cell variability in transient phosphorylation is derived from noise in the intensity of the upstream signals in reaction to the applied external stimulus. During sustained, by contrast, variability stems primarily from noise in the background of the upstream signal as well as in the degradation of the kinase. To further substantiate our results we propose (Silk et al., 2014) and implement new experimental interventions which have allowed us to conclusively rule out any non-negligible impact of intrinsic noise. The workflow adopted in this analysis is summarized in Figure 1E.

**Figure 1:**
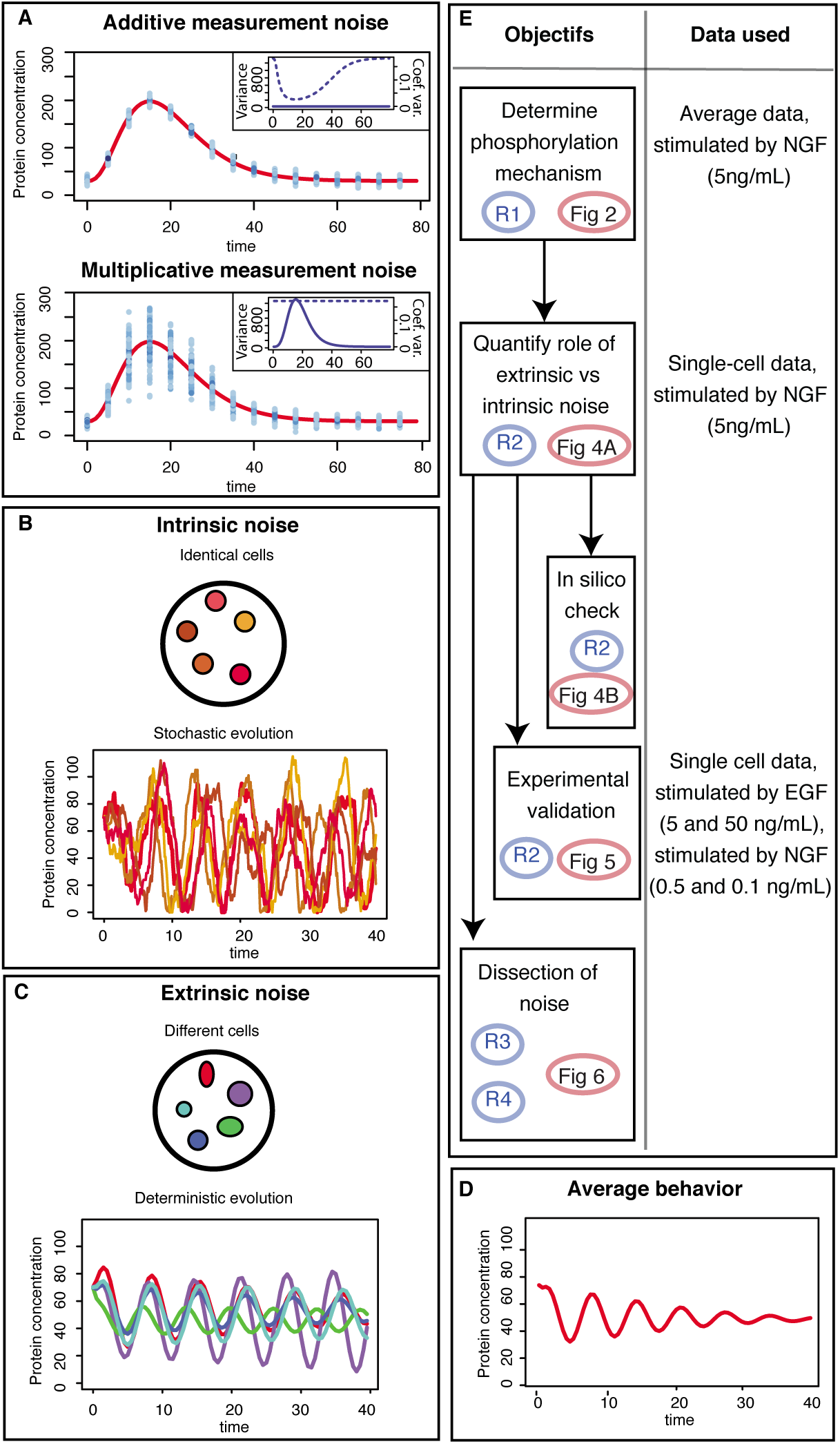
Noise and cell-to-cell variability. **(A)** Effect of additive (top) and multiplicative (bottom) measurement noise. Protein concentrations are shown by the red lines, and samples of 100 noisy data measurements every five minutes are represented by blue dots. In the embedded figures we show the evolution of the variance (solid line) and the coeficient of variation (dotted line). The variance of data with additive measurement noise is constant over time, whereas the variance of data with multiplicative measurement noise varies with time. The opposite behaviour is observed for the coeficient of variation. (B-D) Within-cell variability can be caused by intrinsic noise (B), resulting from the stochastic nature of biochemical reactions, or extrinsic noise (C), arising from inherent differences between the cells. The average evolution of the protein concentration across the heterogeneous cell population is identical whether the cells are subject to intrinsic or extrinsic noise (D). (E) Flow chart of the analysis of origin of cell-to-cell variability as performed in this paper, highlighting which data are used at each steps.The symbols R1 to R4 refer to each of the 4 result subsections and Fig 2 to Fig 6 denotes the figures‥

## Results

### Quantifying temporal evolution of cell-to-cell variability

We investigate the causes of cellular heterogeneity *in vivo* during ERK activation by doubly phosphorylated MEK in PC12 cells. This cell-to-cell variability study is based on measurements of the concentration of doubly phosphorylated MEK and ERK at the single cell level obtained by quantitative image cytometry. Cells are plated in medium containing a fixed amount of neuronal growth factor (NGF) as the stimulus at time *t* = 0. Every two minutes cells in one well are fixed in order to quantify the concentration of the two proteins of interest providing us with a series of cross sectional snapshots of the joint protein distributions of doubly phosphorylated MEK and ERK (i.e the sum of free and complex bound forms), see Figure 3A.

The observed distributions of the total amount of doubly phosphorylated MEK and ERK are illustrated in Figure 3B, and Figure 3C shows the evolution of the variance, the coefficient of variation and the Fano factor over time for both proteins. The variance over the cell population of the concentration is of the order of 10^5^ and significantly varies with time. Because both the coefficient of variation and the variance of the amount of doubly phosphorylated ERK vary with time we can rule out the possibility that the variability in the protein concentration measurements has been caused by additive or multiplicative measurement noise, see Figure 1A. In addition, the experimental noise in QIC has been estimated by Uda et al. (see Figure S2 in (Uda et al., 2013)) and is found to be negligible compared to the level of cell-to-cell variability Any analysis of the origins of cell-to-cell variability requires us to determine the modes of ERK phosphorylation and dephosphorylation. ERK activation involves phosphorylation at both its tyrosine and threonine phosphorylation sites by its cognate kinase MEK (Ferrell and Bhatt, 1997; Ferrell et al., 2014). Previous studies (Toni et al., 2012) have shown that *invivo* phosphorylation (as well as dephosphorylation) occurs in two steps where the kinase binds to the protein twice in order to phosphorylate the two sites successively (see Figure 2B). Using a Bayesian model selection (Kirk et al., 2013) approach, we confirm that this distributive mechanism (Ferrell et al., 2014) best captures the observed average behavior in our data (see *Supplemental Information*). We therefore base our analysis of the origins of cell-to-cell variability on this mechanistic model with 20 model parameters including 12 reaction rates (see Figure 2B Middle and Bottom), 4 parameters describing the impact of the NGF stimulus and upstream signals (see Figure 2B Top) and 4 parameters controlling the initial concentrations of the species involved in the ERK–MEK core system (see *Supplemental Information*).

**Figure 2:**
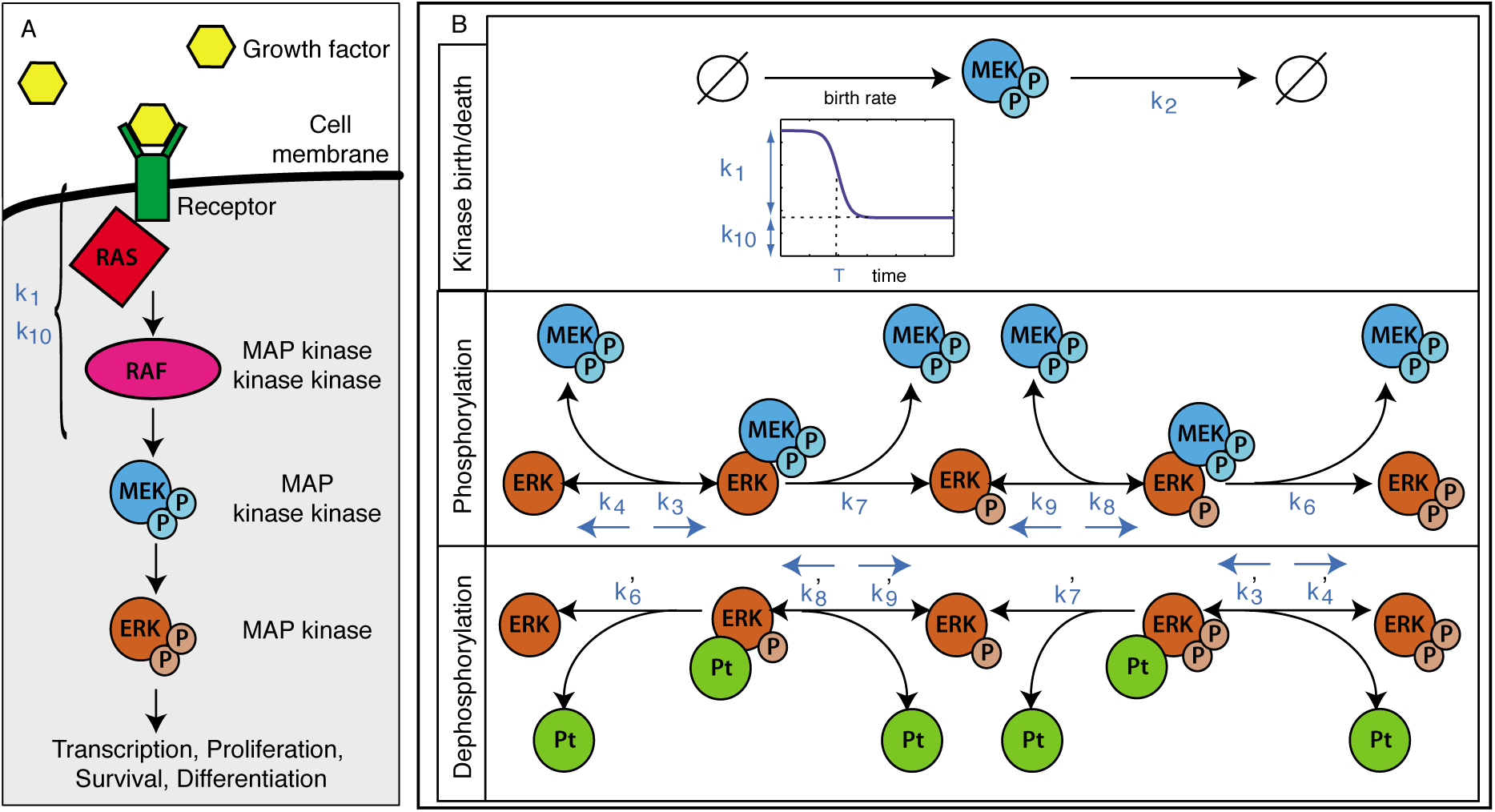
MAPK Signalling. **(A)** The RAS-RAF-ERK signal transduction cascade in response to a neural growth factor (NGF), which activates the membrane-bound GTPase (RAS); this leads to the activation of the RAF kinase and subsequently to the phosphorylation of MEK; active MEK in turn phosphorylates ERK. **(B Top)** The impact of the NGF stimulus and the upstream reactions on the evolution of the concentration of active MEK are modelled using a time dependent function which depends on three parameters: *k*_1_ describes the pulse height, *k*_10_ the background signal and *T* the time at which the influence of the upstream reactions drops down. In addition active MEK is degraded with rate *k*_2_. **(Middle and Bottom)** Mechanistic model describing the phosphorylation and dephosphorylation processes of ERK. Pt denotes the cognate ERK phosphatase. The reaction rates are shown next to their associated reactions.

**Figure 3:**
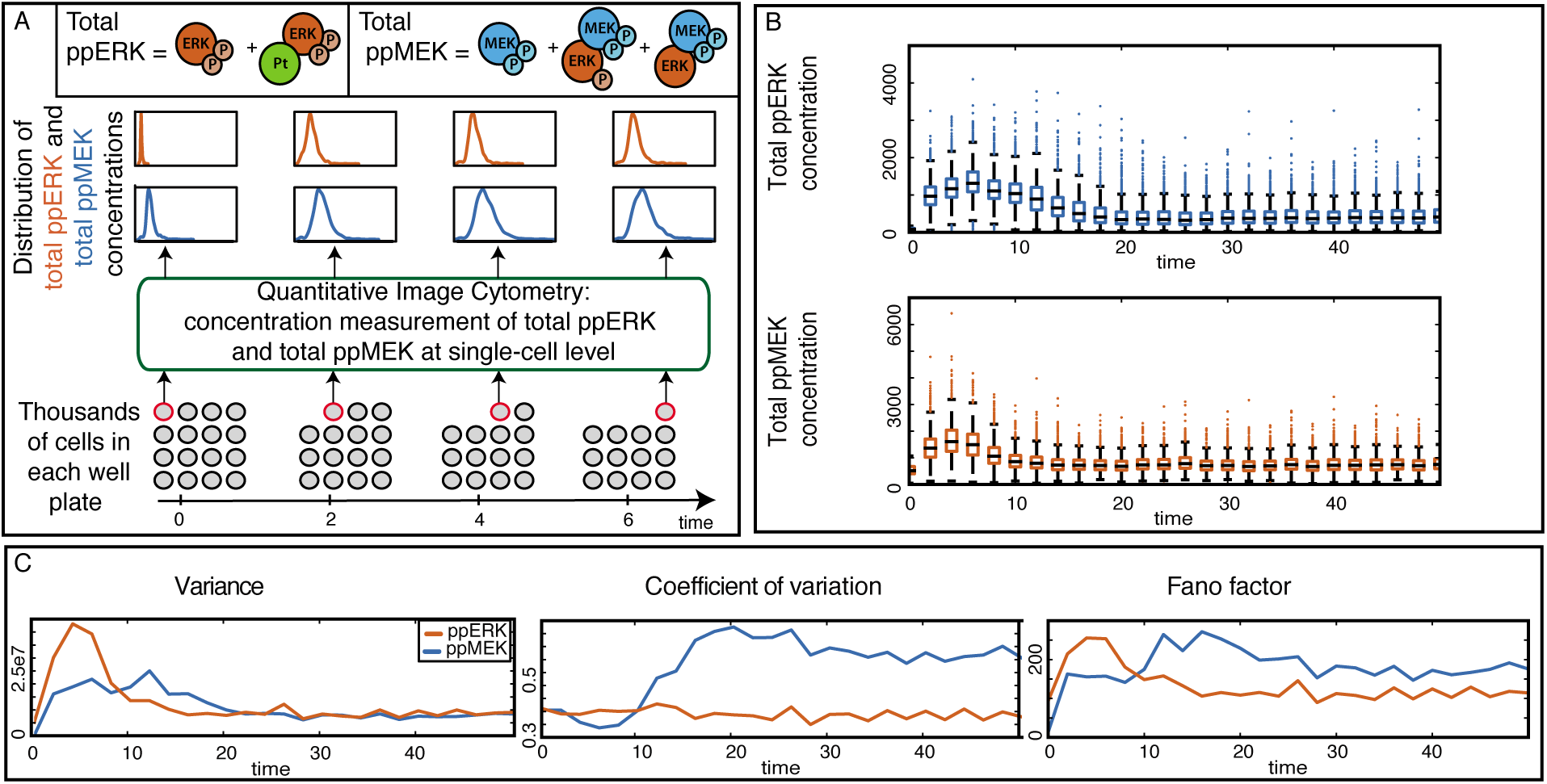
Quantifying temporal evolution of cell-to-cell variability. **(A)** Cells are plated in a medium and stimulated with NGF at *t* = 0. Every two minutes, thousands of cells are fixed and the amount of total doubly phosphorylated MEK and ERK (i.e. the sum of free and complex bound forms) is measured at single-cell level using quantitative image cytometry, providing a series of cross sectional snapshots of the joint distributions of MEK and ERK levels. **(B)** Boxplots showing the distributions of the measured protein concentrations at each time point (the edges of the coloured boxes are the 0.25 and 0.75 quantiles; the central mark is the median). **(C)** The temporal evolution of the variance, the coefficient of variation and the Fano factor for the distributions of the two proteins.

### Intrinsic noise alone cannot explain the observed variabilities between cells

While it is straightforward to model extrinsic and intrinsic noise, quantifying their relative contributions to real molecular systems has thus far only been possible for systems where two-reporter assays are available(Elowitz et al., 2002; Swain et al., 2002). Here we develop a statistical framework that allows us to obtain quantitative insights into the roles of these two sources of noise for signalling systems where direct measurements are typically not possible.

Extrinsic sources of variability stem from all those elements of the “real system” that are not explicitly modelled; these typically include factors such as inherent differences between the cells in terms of cell-size, stage of cell-cycle, protein concentrations at the start of the experiment, and other biophysical parameters. To capture such effects we allow model parameters to differ between cells (Shahrezaei et al., 2008; Toni and Tidor, 2013): the parameters for each cell are drawn from a log-normal distribution (with “hyper-parameters” (Gelman et al., 2013) for means and variances that will be inferred from the data). The potential sources of extrinsic noise are: differences in the reaction rates between cells, different initial concentrations of ERK and MEK, and differences in the upstream signalling cascades feeding into the MEK dynamics.

Using the Bayesian framework developed in *Experimental procedures* and *Supplemental Information* we analyze the roles of intrinsic and extrinsic noise in the single cell data. The resulting statistical model-evidence indicates that the extrinsic noise best explains the data. The evolution of the obtained distributions for MEK and ERK are shown and compared to the data in Figure 4A: only the extrinsic noise model can explain the observed high levels of cell-to-cell variability.

**Figure 4:**
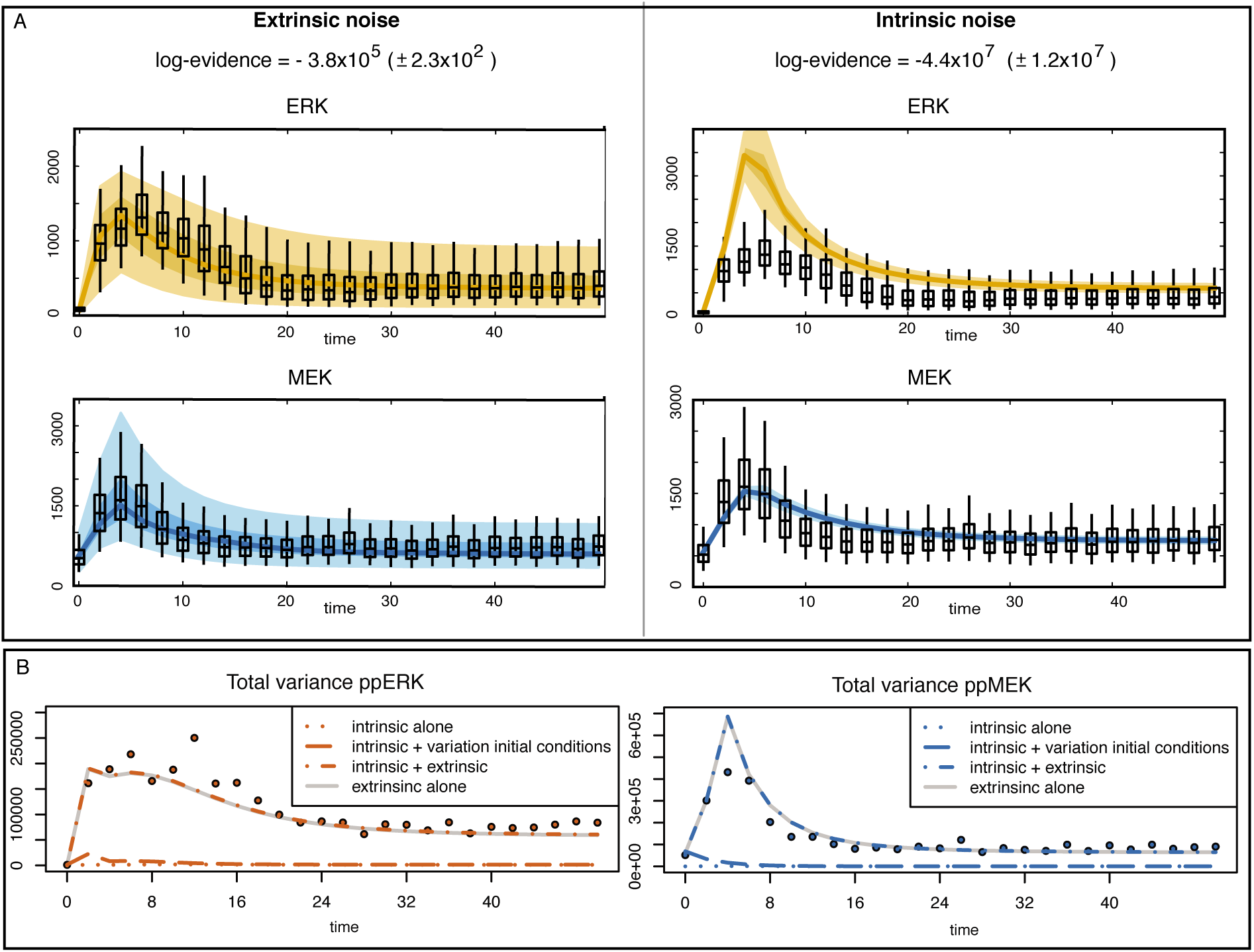
Intrinsic noise alone cannot explain the observed variabilities between cells. (A) Evolution of the inferred distributions over the cell population for the two noise models and comparison to the single-cell data distributions (box plots). The lines represent the median of the distributions, while the shaded regions indicate the regions delimited by 5th and 95th percentiles (lighter regions) and the one delimited by 25th and 75th percentiles (darker regions). The medians and percentiles shown here are the average of the medians and percentiles computed for 1000 sets of parameters sampled from the posterior distribution. The logarithm of the evidence is shown for both noise models; these are strongly supportive of the extrinsic noise models. **(B)** Temporal evolution of the predicted variances over the cell population for the intrinsic noise model alone (dotted lines), the intrinsic noise model combined with a variation in initial conditions between cells (dashed lines), the intrinsic noise together with extrinsic noise (dash-dot lines) and the extrinsic noise alone (grey continuous line), show that the contribution of intrinsic noise is negligible. The dots represent the measured variance of the concentration of the two proteins over the cell populations.

To substantiate this further (and to explore the parameter space more widely) we use Latin hyper-cube sampling to generate a set of 10^6^ parameter vectors and systematically analyse the evolution of the molecular concentrations of MEK and ERK for each of these parameters. Only 20 parameter vectors out of the 10^6^ lead to stable solutions for which the obtained variances of doubly phosphorylated ERK and MEK is higher than 10^5^ (at either 6 or 8 minutes after stimulation; but for none of these parameters do we observe a variance of doubly phosphorylated ERK that is anywhere close to the experimental observations (where the variance is *≈* 3.10^5^).

Variation in initial conditions is also not sufficient to generate the observed cell-to-cell variability; this is easily seen by sampling different values for the the initial concentration of the species involved in the ERK-MEK system according to a log-normal distribution with mean and variance (given by the inferred hyper-parameters for the extrinsic noise case) and simulating the model with intrinsic noise for each of these initial conditions. The total variance, which is the sum of (a) the mean over the different initial conditions of the variance due to the intrinsic noise, and (b) the variance over the different initial conditions of the mean over the intrinsic variability, is shown in Figure 4B. This shows that the variance including variation in initial conditions does not differ appreciably from the variance of intrinsic noise alone.

In a biological system we expect extrinsic and intrinsic sources of noise: the cells are likely to be different in terms of initial molecular concentrations and stage of cell-cycle, and the biochemical reactions occur at random times (Komorowski et al., 2013). We therefore compare the variances of the observed molecular species under extrinsic noise alone with the total variances under both extrinsic and intrinsic noise. From Figure 4B it is apparent that the contribution of intrinsic noise to the total variation is negligible.

An immediate prediction that follows from the above analysis is that the core MEK-ERK system as described here is a reliable and faithful information processing unit: little noise is introduced here, and different signals are mapped onto distinct outcomes in a predictable manner. As a corollary of this we know that the MAPK system does not introduce cell-to-cell variability into the down-stream cellular pathways.

In order to test this prediction and validate the model further we consider the response of the MEK-ERK system to different stimuli; if cell-to-cell variability is due to MEK and ERK dynamics, then the parameterized model developed above should not be able to describe the dynamics. On the contrary, we find that extrinsic noise model can explain the response of the MEK-ERK system to stimulation by EGF, Figure 5A, and different NGF stimulus intensities, Figure 5B (see also also *Supplemental Information* for a more extensive analysis). Here we have used the hyper-parameters inferred previously except for those that correspond to the upstream dynamics (which are known to depend on the stimulus strength and temporal pattern, see (Fujita et al., 2010)); these and only these were inferred directly from the EGF and NGF timecourses. The model with extrinsic noise shows good qualitative and quantitative agreement between model predictions and the new data obtained for different NGF stimulus levels. Thus our extrinsic noise MEK-ERK model is capable of predicting the response to other stimuli than those used in the model development.

**Figure 5:**
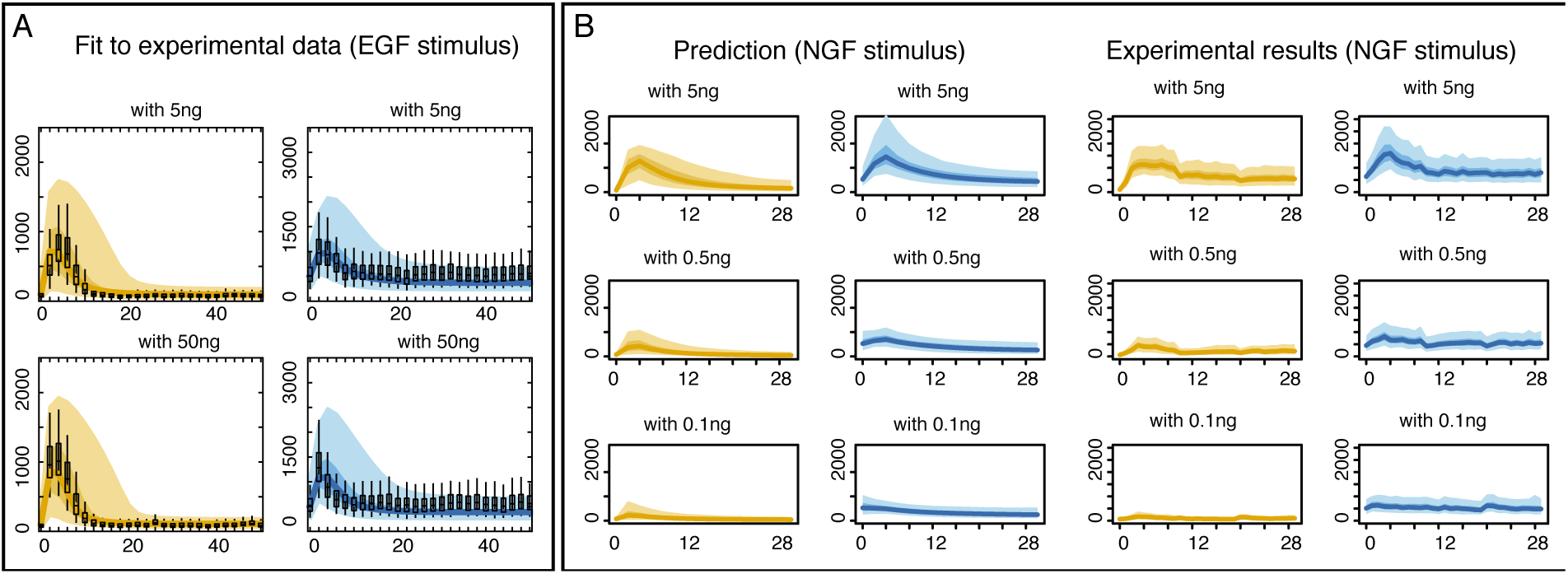
Prediction of the impact of growth factor on cell-to-cell variability. **(A)** Evolution of the inferred distributions of the total amount of doubly phosphorylated ERK and MEK in response to stimulation by EGF with two levels of intensity. The hyper-parameters for the initial conditions and the reaction rates are fixed to the previously estimated values (using the single-cell data in response to NGF stimulus) whereas the hyper-parameters describing the impact of the stimulus and upstream signals on the kinase are inferred here separately. The single-cell data distributions (box plots) are compared to the inferred distributions (the lines represent the median of the distributions, while the shaded regions indicate the regions delimited by 5th and 95th percentiles for the lighter regions and the one delimited by 25th and 75th percentiles for thedarker regions) **(B)** The predictions for the behaviour of the total amount of doubly phosphorylated ERK and MEK under the extrinsic noise model (left columns) are compared to experimental measurements (right columns) for different level of NGF intensity. The solid lines indicate the median value, while the shaded regions indicate the regions delimited by 5th and 95th percentiles (lighter zones) and by 25th and 75th percentiles (darker zones).

### Fluctuations in the upstream reactions and in the degradation rate of the kinase explain most of the cell-to-cell variability

Our Bayesian analysis allows us to assess directly which parameters differ most between cells. For each parameter we have estimates of the *coefficient of variation* across cells, and the parameters that contribute most to the observed cell-to-cell variability are those for which the inferred coefficient of variation is consistently and significantly different from zero (see *Supplemental Information*). We find five strongly contributing factors: three model parameters (*k*_1_, *k*_2_ and *k*_10_) and the two initial conditions that describe the level of background activity present in the cell at the point of stimulation. The pulse height, *k*_1_, and the background upstream signal, *k*_10_, jointly characterise the impact of the NGF stimulus and the upstream reactions on the evolution of active MEK (see Figure 2B) Top. The degradation rate of active MEK (*k*_2_) affects the steady state levels of cell-to-cell variability and the role of degradation reactions in determining levels of noise (and thus cell-to-cell variability) has been well documented (Komorowski et al., 2013). In Figures 6A we illustrate the predominant role that the upstream parameters have on the extent of cell-to-cell variability in this system.

In Figure 6B we further show that that other factors — measurements for cell-size and volume, and Hoechst level, (the dye used to quantify nucleic acid levels) — make only negligible contributions to observed levels of cell-to-cell variability. The total amounts of doubly phosphorylated ERK and MEK have the highest partial correlation and we can thus rule out cell-cycle etc. as explanations for, or cause the temporal variability in the amount of active ERK.

**Figure 6:**
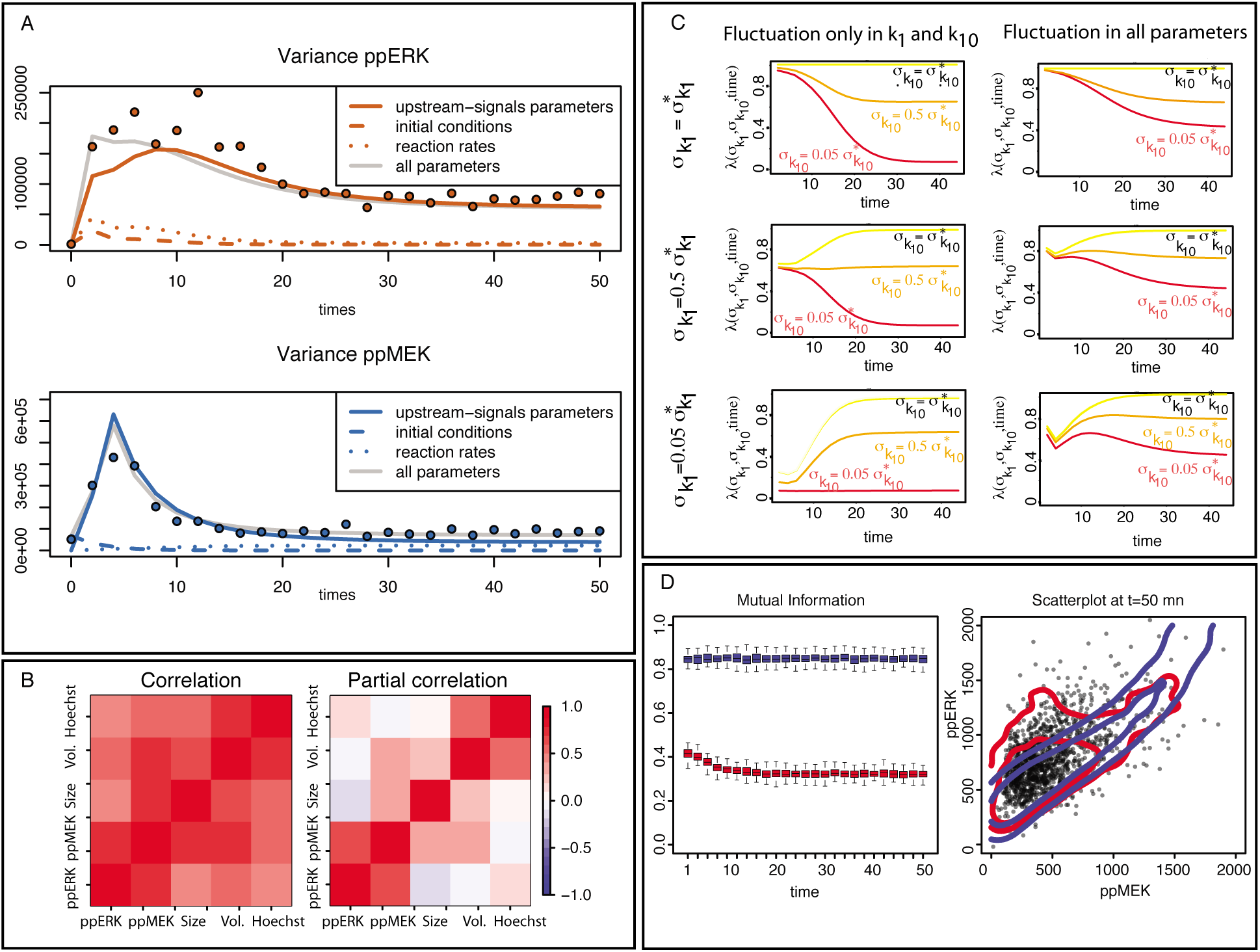
Factors contributing to cell-to-cell variability and its impact on Cellular Information Processing. (A) Evolution of the predicted variances when only some of the model parameters vary from one cell to another: either the parameters that describe the effect of the upstream signals (continuous lines), the parameters controlling the initial conditions (dashed lines) or the reaction rates (dotted lines). The light grey continuous line is the predicted variance when all model parameters differ between cells. The dots represent the measured variance of the concentration of the two proteins over the cell populations. **(B)** Correlation and partial correlation between the measurements of the amount of ppERK and ppMEK, the cell size, the cell volume and the Hoechst intensity (both plots are on the same scale, see colour bar). **(C)** Impact of the variability of the input (i.e. the upstream reactions), described by the variances 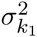 and 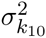 on the level of cell-to-cell variability of the system’s output (i.e. ppERK) which is quantified by the function *λ*(*σ*_*k*1_, *σ*_*k*10_, *t*). *λ*(*σ*_*k*1_, *σ*_*k*10_, *t*) is close to 1 (resp. 0) if the level of cell-to-cell variability in the system’s output is equal (resp. very different) to the level of cellto-cell variability when *σ*_*k*1_ and *σ*_*k*10_ are maximal (i.e equal to *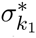* and 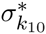). This subfigure is divided in two parts: in the left column, only parameters *k*_1_ and *k*_10_ vary between cells (all other model parameters are fixed to their mean value), whereas in the right column the full extrinsic noise model is considered where all model parameters differ between cells. For each column, the behaviour of *λ*(*σ*_*k*1_, *σ*_*k*10_, *t*) with time is illustrated for decreasing values of *σ*_*k*1_ and *σ*_*k*10_. Each panel corresponds to a fixed value of *σ*_*k*1_ while each line corresponds to a fixed value of *σ*_*k*10_, the red corresponding to the smallest and the yellow to the maximum standard deviation. **(D)** Mutual information between ppERK and ppMEK (Left) when the system is simulated under the full extrinsic noise model (red) or only varying the parameters *k*_1_ and *k*_10_ between cells (blue). Comparison of the joint distribution of the concentration of ppERK and ppMEK when the system is simulated under both models (blue and red) as well as the joint distribution of the data (black dots).

### Impact of cell-to-cell variability on Cellular Information Processing

We conclude our analysis by investigating the role that noise plays in mediating the response of MAPK signalling cascades to external stimuli. We analyse the level of cell-to-cell variability in the system’s output (i.e. the total amount of doubly phosphorylated ERK) as a function of how variable the inputs (captured by the transient and sustained upstream intensities, *k*_1_ and *k*_10_, and their respective variances over the cell population 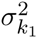 and 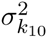 are. We simulate system output for given values of 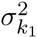 and 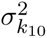 and compute the ratio

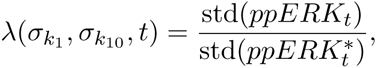

where std(*ppERK*_*t*_) is the standard deviation of the output at time *t*, and std(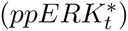) is the standard deviation of the system’s output at time *t* if the variance of the input is maximal (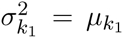 and 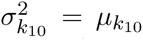 where *μ*_*k*1_ and *μ*_*k*10_ are the means over the cell population for, respectively, *k*_1_ and *k*_10_). This ratio quantifies the change in the level of cell-to-cell variability in the system’s output as the input noise is decreased.

In the first instance we assume that only the input signal strengths (*k*_1_ and *k*_10_) vary between cells — all other model parameters are fixed to the inferred posterior mean values. The evolution of *λ*(*σ*_*k*1_, *σ*_*k*10_, *t*) over time when varying the variances 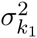 and 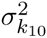 is shown in Figure 6C (left column). Before *t* = 8 minutes, *λ*(*σ*_*k*1_, *σ*_*k*10_, *t*) increases with 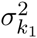 whereas 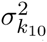 has no impact on *λ*(*σ*_*k*1_, *σ*_*k*10_, *t*). Conversely, after *t* = 24 minutes, *λ*(*σ*_*k*1_, *σ*_*k*10_, *t*) increases with *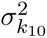* but 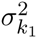 no longer affects output variability. Thus variability in active ERK abundance across the cell population is initially strongly influenced by the variability in pulse height, and subsequently by the variability in the sustained or background signal.

To investigate the effect of the variability in all model parameters on cellular information processing, we also simulate the system under extrinsic noise (varying all model parameters between cells), and compute once more *λ*(*σ*_*k*1_, *σ*_*k*10_, *t*) for different signal variabilities. It is apparent from Figure 6C (right column) that, under the extrinsic noise model, the level of cell-to-cell variability in the system’s output remains substantially high even when the variability in the system’s input has been decreased considerably (*λ ∽* 0.45 when *σ*_*k*1_ and *σ*_*k*10_ are divided by 20). Therefore, the presence of extrinsic noise weakens the influence of the variability in the upstream signal upon the cell-to-cell variability in the system’s output.

To follow on from this, we compute the mutual information between the total amount of ppMEK and the total amount of ppERK at different time points, simulating the system under extrinsic noise or varying only the parameters that seems to be related to most of the cellular variability (*k*_1_, *k*_2_ and *k*_10_). We observe in Figure 6D (Left) that the presence of extrinsic noise decreases the level of transfer of information between the two species of interest. This difference can be easily explained by comparing the joint distribution of the concentration of ppERK and ppMEK when the system is simulated under the full extrinsic noise model or only varying the ‘driving’ parameters (see Figure 6D Right). Even though only varying the ‘driving’ parameters explain the evolution of the variance and correlation between the two proteins, only the full extrinsic noise model captures the shape of the joint distribution.

## Discussion

We have used quantitative image cytometry to elucidate the causes of population heterogeneity in the MAPK signaling cascade and presented a comprehensive analysis of cell-to-cell variability in the activation dynamics of the MEK–ERK system to environmental stimuli. Our analysis shows that the *in vivo* modes of ERK phosphorylation and dephosphorylation are distributive. With a reliable model for the (de–)phosphorylation mechanisms(Toni et al., 2012) in hand, we were then able to dissect the nature of the cell-to-cell variability inherent in the data. Recent MAPK models proposed in the literature (Ferrell et al., 2014; Harrington et al., 2013; Ortega et al., 2006; Sturm et al., 2010; Voliotis et al., 2014) allow for very rich dynamics and *a priori* it is therefore impossible to make an appeal to the large number of MEK, ERK and other molecules present in the eukaryotic cell, in order to rule out a role for intrinsic noise.

The detailed analysis of these alternative mechanisms gives a clear verdict in favour of extrinsic noise as the dominant factor for the observed cell-to-cell variability in the MEK–ERK system. Few, if any parameters appear to be tightly constrained across the populations of cells considered here. For some parameters we do, in fact, find strong evidence that they vary quite considerably between cells; but the MEK–ERK core system itself adds little to the observed levels of cell-to-cell variability, and is capable of transmitting upstream information faithfully. Thus differences in reactions upstream from the MEK–ERK core are passed on by the cascade to the downstream machinery. We propose that cells employ temporal selection of different noise sources for their intra-cellular information processing. In particular, we show that the cell-to-cell variability during sustained phosphorylation stems from random fluctuations in the background or base-line upstream signalling processes, while during transient phosphorylation, the cellular heterogeneity in ERK activity is primarily due to noise in the intensity of the upstream signal(s). The stage at which a cell is in its cell cycle is an obvious potential cause for cell-to-cell variability, but here we find that this can explain only a fraction of the overall extent of heterogeneity in the abundance of active ERK.

We found that extrinsic noise in the MAPK system considered here tends to attenuate variability in the up-stream signal prior to it arriving at MEK. The distributively operating MEK–ERK systems is furthermore capable of kinetic proof-reading (Hlavacek et al., 2001; Murugan et al., 2012), and the combination of this mechanism with the behaviour observed for the extrinsic noise, makes this a very effective filter for noisy upstream signals, especially at the population-level. Given the importance of MAPK systems in different cell-fate decision making processes such robustness to noise is clearly important. But while kinetic proof-reading confers robustness to all cells similarly, the extrinsic variability will mean that some cells may be better poised to process environmental signals subject to noise than others, which would lend robustness at the population-level, similar to bet-hedging behaviour in evolutionary biology (Kussell and Leibler, 2005; Stumpf et al., 2002). In development and tissue homeostasis (Rué and Arias-Martinez, 2015) (and in regenerative medicine) it may be important to find ways to regulate population-level behaviour further and here other, inter- and intra-cellular feedback mechanisms that control cell-to-cell variability further (Michailovici et al., 2014).

The study presented here is based on experiments carried out in PC12 cell lines(Greene and Tischler, 1976), which unlike *in vitro* set-ups, provide the cell physiological context. The activity of up-stream and down-stream processes affecting ERK may depend on cell-type; this has, for example, been shown for nuclear shuttling, where even subtle differences between different cell lines can affect e.g. the activity of nuclear ERK (Harrington et al., 2012). Our deliberate focus on the core MEK–ERK dynamics is less prone to such strong cell-type specificity over the time-scales considered, whereas the potential of feedback from either ERK or any of its many down-stream targets onto the MAPK cascade or proteins further upstream should be carefully considered in different cell-types. The additional richness in behaviour that such feedback (Ortega et al., 2006; Sturm et al., 2010) or explicit consideration of nuclear shuttling (Harrington et al., 2013; Mugler et al., 2013) of ERK and MEK can induce warrants further investigation (Ozaki et al., 2010); here over the time-course considered, and in light of the data available such effects are marginal, but this may change as other or longer stimuli, or more complex temporal stimulation patterns are considered. At single cell level both feedback and shuttling — the latter especially if it induces multi-stability — are therefore clearly worth of further investigation; there, however, we may also have to consider differences between cell-lines or cell types (Harrington et al., 2012). It is important to keep in mind that no model will ever be able to contain all the constituent parts of any biological system of any real-world relevance (Babtie et al., 2014). Therefore extrinsic noise — variation due to factors not explicitly included in the model — will always be an issue for modelling molecular and cellular systems. The present work shows that this need not necessarily limit the usefulness or usability of mechanistic, mathematical models of biological systems. By pinpointing the sources of extrinsic noise, which are typically not obvious *a priori*, sound statistical modelling is able to provide deeper mechanistic insights and highlight where a model ought to be extended, or whether this is indeed necessary.

## Experimental procedures

### Experimental data collecting process

The concentrations of molecular species were measured using quantitative image cytometry (QIC) (Ozaki et al., 2010; Saito et al., 2013). PC12 cells were seeded at a density of 10^4^ cells per well in 96-well poly-L-lysinecoated glass-bottomed plates (Thermo Fisher Scientific, Pittsburgh, PA). 24 hours after seeding, the medium was replaced with DMEM containing 25mM HEPES and 0.1 percent of bovine serum albumin. 18 hours after serum starvation, the stimulus is applied by replacing the starvation serum with a medium containing the stimulant (5 or 0.5 or 0.1ng/mL). Our setup carries out stimulation in an incubator and achieved 1-minute interval stimulation at 37^°^C under 5% CO2 in saturated air humidity. The cells are then fixed with 4 percent paraformaldehyde for 10 minutes and immunostained. Cells were subjected to QIC analysis with mouse anti- ppERK Sigma Aldrich M8159 antibody and rabbit anti–pMEK Cell Signaling Technology 9121. Note that anti–pMEK antibody detects both singly and doubly phosphorylated MEK.

All images were analyzed with Cell Profiler (Kamentsky et al., 2011). The nuclear region was identified based on Hoechst imaging, and the cellular region was identified based on CellMask stained images going out from from the nuclear region. Total cellular signal intensity in nuclear regions and cellular regions were measured for ppERK and pMEK, respectively. We used the cellular region in pixels as the cell size and the intensity of CellMask in the cellular region as a measure of cell volume.

### Parameter inference and model evidence

We use a Bayesian approach in order to infer the parameters of the system (see *Supplemental Information* for a detailed list of the model parameters) and rank the candidate mechanistic models. Bayesian parameter inference is centred around the posterior probability distribution, 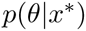, which strikes a compromise between prior knowledge, *p*(*θ*), about parameter vectors, *θ*, and the capacity of a parameter to explain the observed data, *x*^*\**^, measured by the likelihood *p*(*x*^*\**^*|θ*), via

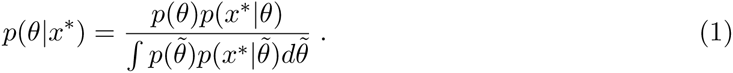

Here we evaluate the posterior using a sequential Monte Carlo (SMC) sampler proposed by (Del Moral et al., 2006), which is easily parallelized. The output of the algorithm is a set of weighted parameter vectors *{θ*^(*i*)^, *ω*^(*i*)^*}*_1*≤i≤N*_. Here the parameter vector associated to the highest weight is called the *inferred parameter vector*. Technical details about our implementation of the SMC sampler algorithm are given in the *Supplemental Information*.

The SMC sampler algorithm also enables us to compute the *model evidence* (Kirk et al., 2013), which is the probability to observe the data *x*^*\**^ under the model *ℳ* (given the alternative models considered),

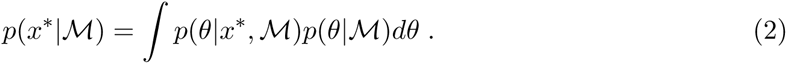

The model evidence allows us to rank candidate models in terms of their ability to explain the observed data *x*^*\**^: the best model is the one with the highest model evidence. In addition, the Bayes factor assesses the plausibility of two candidate models *ℳ*_1_ and *ℳ*_2_:

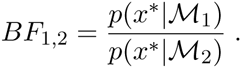

Whenever *BF*_1,2_ is larger than 30, the evidence in favour of model *ℳ*_1_ is considered very strong (Jeffreys, 1961). We use our own implementation of the SMC sampler algorithm in Python as well as an interface to simulate the models in a computational efficient manner using a GPU accelerated ODE solver (Zhou et al., 2011) and a C++ ODE solver for stiff models (Hindmarsh et al., 2005).

### Likelihood functions

At each time point *t ∈* 𝕋 = *{*0, 2, 4, *…*50*}* the concentrations of the pMEK and ppERK are measured in *N*_*t*_ different cells. We denote by 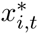 and 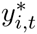 the concentration of the two proteins in the *i*-th cell, 1 *≤ i ≤ N*_*t*_, and by 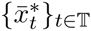 and 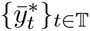 the observed average trajectories. In addition we denote by *x*_*t*_(*θ*) and *y*_*t*_(*θ*) the solution of the system of ODE given the parameter vector *θ* at time *t*.

Assuming an independent Gaussian measurement error for each time point with constant variance *v*, the likelihood function for the average data measurements is

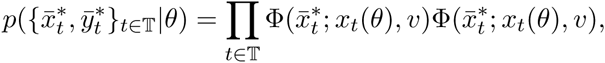

where Φ(*·*; *m, v*) is the probability density function of a normal distribution of mean *m* and variance *v*. The variance *v* is inferred simultaneously with the other parameters.

In order to derive the likelihood function in the intrinsic noise model we use the linear noise approximation (LNA). The LNA provides a system of ODEs which describe how the means and the variances of the molecular species vary over time. These equations are produced using the *StochSens* package (Komorowski et al., 2012). With *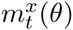,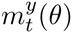,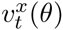* and 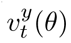 denoting the solutions of the ODEs describing the means and variances for the parameter *θ* at time *t*, the likelihood 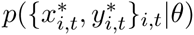 is equal to

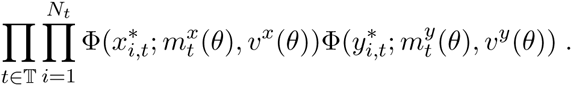

Extrinsic noise is modelled by considering that each cell has a different set of parameters. The distribution of each parameter across the cell population is assumed to be log-normal. We assume that these distributions are independent and denote by *μ*_*θ*_ and 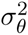 the vector of the means and variances of these distribution, respectively. There is no closed-form expression for the probability *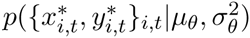* and we use the so-called Unscented Transform (UT), which, given the first two moments *μ*_*θ*_ and 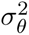 of the distribution in the parameter space, provides an approximation of the evolution of the means and variances of the two species of interest. We denote by 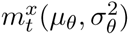 and 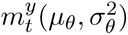 the resulting mean behaviours of the two species at time *t*, and by 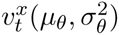 and 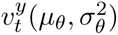 the associated variances. Assuming that the concentration of the doubly phosphorylated ERK and MEK proteins are log-normally distributed we obtain that the likelihood 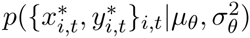 is

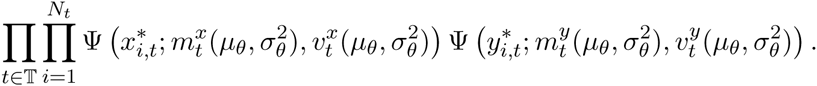

Here Ψ(*·*; *m, v*) is the probability density function of a log–normal distribution with mean *m* and variance *v*. The *Supplemental Information* contains additional technical details on the computation and the UT algorithm.

### Latin Hypercube sampling

We use Latin Hypercube sampling (LHS) (McKay et al., 1979) to generate 10^6^ parameter vectors in a 20-dimensional space using the Matlab function *lhsdesign*.

### Correlation analysis

In addition to the experimental measurements for the total amount of doubly phosphorylated ERK and MEK our assay also obtained measurements for cell-size, cell volume and Hoechst intensity in each cell. We computed the correlation and partial correlations between these 5 measurements using the R package *GeneNet* (Schäfer et al., 2001).

### Mutual information

The mutual information between two species (ppERK and ppMEK) is computed based on measurements of the protein concentrations in single-cells at different time points. For each time point, we estimate the mutual information using a kernel density estimate of the joint distribution. We use a gaussian kernel with a diagonal covariance matrix and marginal variances equal to 1.06*σN*^-1/5^ where *σ* is the marginal variance of the data and *N* is the number of data points (Silverman, 1986).

## Author contributions

SF, CPB, SK and MPHS designed the study; TK, TT and SK performed the data collection and initial processing; SF, CPB, PK performed modelling and statistical analysis; SF, CPB, PK, SK and MPHS wrote the paper; all authors approved the final version of the manuscript.

## Acknowledgements

PK, TK, SK and MPHS acknowledge financial support from the *Human Frontiers Science Programme*; SF is funded through an MRC Biocomputing fellowship; CPB is a Wellcome Trust Career Development Fellow; SF, CPB, PK, SK and MPHS were also funded through a JST/BBSRC partnering award. MPHS is Royal Society Wolfson Research Merit Award holder.

## References

Aoki, K., Yamada, M., Kunida, K., Yasuda, S. and Matsuda, M. 2011. Processive phosphorylation of ERK MAP kinase in mammalian cells. Proceedings of the National Academy of Sciences, USA 108, 12675–12680.

Ashall, L., Horton, C. A., Nelson, D. E., Paszek, P., Harper, C. V., Sillitoe, K., Ryan, S., Spiller, D. G., Unitt, J. F., Broomhead, D. S., Kell, D. B., Rand, D. A., Sée, V. and White, M. R. H. 2009. Pulsatile stimulation determines timing and specificity of NF-kappaB-dependent transcription. Science 324, 242–246.

Babtie, A. C., Kirk, P. and Stumpf, M. P. 2014. Topological sensitivity analysis for systems biology. Proceedings of the National Academy of Sciences of the United States of America 111, 18507–18512.

Balázsi, G., van Oudenaarden, A. and Collins, J. J. 2011. Cellular decision making and biological noise: from microbes to mammals. Cell 144, 910–925.

Batchelor, E., Loewer, A., Mock, C. and Lahav, G. 2011. Stimulus-dependent dynamics of p53 in single cells. Molecular Systems Biology 7, 488.

Bowsher, C. G. and Swain, P. S. (2012a). Identifying sources of variation and the flow of information in biochemical networks. Proceedings of the National Academy of Sciences 109, E1320–E1328.

Bowsher, C. G. and Swain, P. S. (2012b). Identifying sources of variation and the flow of information in biochemical networks. Proceedings of the National Academy of Sciences, USA 109, E1320–E1328.

Colman-Lerner, A., Gordon, A., Serra, E., Chin, T., Resnekov, O., Endy, D., Pesce, C. G. and Brent, R. 2005. Regulated cell-to-cell variation in a cell-fate decision system. Nature 437, 699–706.

Del Moral, P., Doucet, A. and Jasra, A. 2006. Sequential monte carlo samplers. Journal of the Royal Statistical Society: Series B (Statistical Methodology) 68, 411–436.

Elowitz, M. B., Levine, A. J., Siggia, E. D. and Swain, P. S. 2002. Stochastic gene expression in a single cell. Science 297, 1183–1186.

Ferrell, J. E. and Bhatt, R. R. 1997. Mechanistic studies of the dual phosphorylation of mitogen-activated protein kinase. Journal of Biological Chemistry 272, 19008–19016.

Ferrell, J. E., Jr and Ha, S. H. 2014. Ultrasensitivity part II: multisite phosphorylation, stoichiometric inhibitors, and positive feedback. Trends in Biochemical Sciences 39, 556–569.

Fujita, K. A., Toyoshima, Y., Uda, S., Ozaki, Y.-i., Kubota, H. and Kuroda, S. 2010. Decoupling of receptor and downstream signals in the Akt pathway by its low-pass filter characteristics. Science Signaling 3, ra56–ra56.

Gelman, A., Carlin, J. B., Stern, H. S., Dunson, D. B., Vehtari, A. and Rubin, D. B. 2013. Bayesian data analysis. CRC press.

Geva-Zatorsky, N., Rosenfeld, N., Itzkovitz, S., Milo, R., Sigal, A., Dekel, E., Yarnitzky, T., Liron, Y., Polak, P., Lahav, G. and Alon, U. 2006. Oscillations and variability in the p53 system. Molecular Systems Biology 2, 2006.0033.

Greene, L. A. and Tischler, A. S. 1976. Establishment of a noradrenergic clonal line of rat adrenal pheochromocytoma cells which respond to nerve growth factor. Proceedings of the National Academy of Sciences 73, 2424–2428.

Harrington, H. A., Feliu, E., Wiuf, C. and Stumpf, M. M. P. 2013. Cellular compartments cause multistability in biochemical reaction networks and allow cells to process more information. Biophysical Journal 104, 1824–1831.

Harrington, H. A., Komorowski, M., Beguerisse-Diaz, M., Ratto, G. M. and Stumpf, M. P. H. 2012. Mathematical modeling reveals the functional implications of the different nuclear shuttling rates of Erk1 and Erk2. Physical Biology 9, 036001.

Hilfinger, A. and Paulsson, J. 2011. Separating intrinsic from extrinsic fluctuations in dynamic biological systems. Proceedings of the National Academy of Sciences, USA 108, 12167–12172.

Hindmarsh, A. C., Brown, P. N., Grant, K. E., Lee, S. L., Serban, R., Shumaker, D. E. and Woodward, C. S. 2005. SUNDIALS: Suite of nonlinear and differential/algebraic equation solvers. ACM Transactions on Mathematical Software (TOMS) 31, 363–396.

Hlavacek, W. S., Redondo, A., Metzger, H., Wofsy, C. and Goldstein, B. 2001. Kinetic proof-reading models for cell signaling predict ways to escape kinetic proofreading. Proceedings of the National Academy of Sciences 98, 7295–7300.

Jeffreys, H. 1961. Theory of Probability. 3 edition, Oxford University Press, USA.

Jeschke, M., Baumgartner, S. and Legewie, S. 2013. Determinants of Cell-to-Cell Variability in Protein Kinase Signaling. PLOS Computational Biology 9.

Kamentsky, L., Jones, T. R., Fraser, A., Bray, M.-A., Logan, D. J., Madden, K. L., Ljosa, V., Rueden, C., Eliceiri, K. W. and Carpenter, A. E. 2011. Improved structure, function and compatibility for CellProfiler: modular high-throughput image analysis software. Bioinformatics 27, 1179–1180.

Kiel, C. and Serrano, L. 2009. Cell Type-Specific Importance of Ras-c-Raf Complex Association Rate Constants for MAPK Signaling. Science Signaling 2, ra38–ra38.

Kirk, P., Thorne, T. and Stumpf, M. P. 2013. Model selection in systems and synthetic biology. Current Opinion in Biotechnology 24, 767–774.

Komorowski, M., Finkenstädt, B. and Rand, D. 2010. Using a single fluorescent reporter gene to infer half-life of extrinsic noise and other parameters of gene expression. Biophysical Journal 98, 2759–2769.

Komorowski, M., Miekisz, J. and Stumpf, M. P. 2013. Decomposing Noise in Biochemical Signalling Systems Highlights the Role of Protein Degradation. Biophysical Journal 104, 1783–1793.

Komorowski, M., Žurauskienė, J. and Stumpf, M. P. 2012. StochSensmatlab package for sensitivity analysis of stochastic chemical systems. Bioinformatics 28, 731–733.

Kussell, E. and Leibler, S. 2005. Phenotypic diversity, population growth, and information in fluctuating environments. Science 309, 2075–2078.

McKay, M. D., Beckman, R. J. and Conover, W. J. 1979. Comparison of three methods for selecting values of input variables in the analysis of output from a computer code. Technometrics 21, 239–245.

Michailovici, I., Harrington, H. A., Azogui, H. H., Yahalom-Ronen, Y., Plotnikov, A., Ching, S., Stumpf, M. P., Stumpf, M. P. H., Klein, O. D., Seger, R. and Tzahor, E. 2014. Nuclear to cytoplasmic shuttling of ERK promotes differentiation of muscle stem/progenitor cells. Development 141, 2611–2620.

Mody, A., Weiner, J. and Ramanathan, S. 2009. Modularity of MAP kinases allows deformation of their signalling pathways. Nature cell biology 11, 484–491.

Mugler, A., Tostevin, F. and ten Wolde, P. R. 2013. Spatial partitioning improves the reliability of biochemical signaling. Proceedings of the National Academy of Sciences, USA 110, 5927–5932.

Murugan, A., Huse, D. A. and Leibler, S. 2012. Speed, dissipation, and error in kinetic proofreading. Proceedings of the National Academy of Sciences 109, 12034–12039.

Nelson, D. E., Ihekwaba, A. E. C., Elliott, M., Johnson, J. R., Gibney, C. A., Foreman, B. E., Nelson, G., See, V., Horton, C. A., Spiller, D. G., Edwards, S. W., McDowell, H. P., Unitt, J. F., Sullivan, E., Grimley, R., Benson, N., Broomhead, D., Kell, D. B. and White, M. R. H. 2004. Oscillations in NF-kappaB signaling control the dynamics of gene expression. Science 306, 704–708.

Ortega, F., Garcés, J. L., Mas, F., Kholodenko, B. N. and Cascante, M. 2006. Bistability from double phosphorylation in signal transduction. Kinetic and structural requirements. The FEBS journal 273, 3915–3926.

Ozaki, Y.-i., Uda, S., Saito, T. H., Chung, J., Kubota, H. and Kuroda, S. 2010. A quantitative image cytometry technique for time series or population analyses of signaling networks. PLOS one 5, e9955.

Paszek, P., Ryan, S., Ashall, L., Sillitoe, K., Harper, C. V., Spiller, D. G., Rand, D. A. and White, M. R. H. 2010. Population robustness arising from cellular heterogeneity. Proceedings of the National Academy of Sciences, USA 107, 11644–11649.

Piala, A. T., Humphreys, J. M. and Goldsmith, E. J. 2014. MAP Kinase Modules: The Excursion Model and the Steps that Count. Biophysical journal 107, 2006–2015.

Pujadas, E. and Feinberg, A. P. 2012. Regulated noise in the epigenetic landscape of development and disease. Cell 148, 1123–1131.

Quaranta, V. and Garbett, S. P. 2010. Not all noise is waste. Nature methods 7, 269–272.

Rué, P. and Arias-Martinez, A. 2015. Cell dynamics and gene expression control in tissue homeostasis and development. Molecular Systems Biology 11, 792–792.

Saito, T. H., Uda, S., Tsuchiya, T., Ozaki, Y.-i. and Kuroda, S. 2013. Temporal Decoding of MAP Kinase and CREB Phosphorylation by Selective Immediate Early Gene Expression. PLOS one 8, e57037.

Schäfer, J., Opgen-Rhein, R. and Strimmer, K. 2001. Reverse engineering genetic networks using the GeneNet package. Journal of the American Statistical Association 96, 1151–1160.

Selimkhanov, J., Taylor, B., Yao, J., Pilko, A., Albeck, J., Hoffmann, A., Tsimring, L. and Wollman, R. 2014. Accurate information transmission through dynamic biochemical signaling networks. Science 346, 1370–1373.

Shahrezaei, V., Ollivier, J. F. and Swain, P. S. 2008. Colored extrinsic fluctuations and stochastic gene expression. Molecular systems biology 4.

Silk, D., Kirk, P., Barnes, C. P., Toni, T. and Stumpf, M. P. H. 2014. Model Selection in Systems Biology Depends on Experimental Design. PLOS computational biology 10, e1003650.

Silverman, B. W. 1986. Density estimation for statistics and data analysis, vol. 26,. CRC press.

Spencer, S. L., Gaudet, S., Albeck, J. G., Burke, J. M. and Sorger, P. K. 2009. Non-genetic origins of cell-to-cell variability in TRAIL-induced apoptosis. Nature 459, 428–432.

Spiller, D. G., Wood, C. D., Rand, D. A. and White, M. R. H. 2010. Measurement of single-cell dynamics. Nature 465, 736–745.

Stumpf, M. P. H., Laidlaw, Z. and Jansen, V. A. A. 2002. Herpes viruses hedge their bets. Proceedings of the National Academy of Sciences 99, 15234–15237.

Sturm, O. E., Orton, R., Grindlay, J., Birtwistle, M., Vyshemirsky, V., Gilbert, D., Calder, M., Pitt, A., Kholodenko, B. and Kolch, W. 2010. The mammalian MAPK/ERK pathway exhibits properties of a negative feedback amplifier. Science Signaling 3, ra90.

Swain, P. S., Elowitz, M. B. and Siggia, E. D. 2002. Intrinsic and extrinsic contributions to stochasticity in gene expression. Proceedings of the National Academy of Sciences of the United States of America 99, 12795–12800.

Takahashi, K., Tanase-Nicola, S. and ten Wolde, P. R. 2010. Spatio-temporal correlations can drastically change the response of a MAPK pathway. Proceedings of the National Academy of Sciences, USA 107, 2473–2478.

Toni, T., Ozaki, Y.-i., Kirk, P., Kuroda, S. and Stumpf, M. P. H. 2012. Elucidating the in vivo phosphorylation dynamics of the ERK MAP kinase using quantitative proteomics data and Bayesian model selection. Molecular BioSystems 8, 1921–1929.

Toni, T. and Tidor, B. 2013. Combined model of intrinsic and extrinsic variability for computational network design with application to synthetic biology. PLOS Computational Biology 9, e1002960.

Uda, S., Saito, T. H., Kudo, T., Kokaji, T., Tsuchiya, T., Kubota, H., Komori, Y., Ozaki, Y.-i. and Kuroda, S. 2013. Robustness and compensation of information transmission of signaling pathways. Science 341, 558–561.

Voliotis, M., Perrett, R. M., McWilliams, C., McArdle, C. A. and Bowsher, C. G. 2014. Information transfer by leaky, heterogeneous, protein kinase signaling systems. Proceedings of the National Academy of Sciences 111, E326–E333.

Zhou, Y., Liepe, J., Sheng, X., Stumpf, M. P. and Barnes, C. 2011. GPU accelerated biochemical network simulation. Bioinformatics 27, 874–876.

